# Sex-specific dichotomy of chronic mild stress effects on blood pressure and longitudinal measurements of renal sympathetic nerve activity: are females really protected?

**DOI:** 10.64898/2026.08.04.742819

**Authors:** Dragana Komnenov, Yaaqub Uthman, Noah Ramirez, Christopher T. Banek

## Abstract

Modulation of renal nerves to improve blood pressure (BP) control has become a topic of intense investigation over the last 10-15 years. Given that renal innervation is composed of mixed nerve fibers containing both afferent (sensory) and efferent (sympathetic) fibers, subsequent preclinical studies have been investigating their respective roles in hypertension pathobiology in different genetic and salt-sensitive rat models. Here we set out to investigate how renal afferent and efferent nerves regulate hypertension development in the chronic mild stress model (CMS). We show that in male CMS rats, ablation of afferent renal nerves (ARDNx) and all renal nerves (TRDNx) resulted in similar BP (104 ± 2 mmHg vs. 101 ± 3 mmHg, respectively), both reduced compared to the SHAM group (118 ± 1 mmHg, p = 0.003 and p < 0.001, respectively) arguing for a prominent role of afferent renal nerves in CMS hypertension. Additionally, we show a reduction of vasopressin (AVP) V1b but not V1a receptor abundance in ARDNx CMS males but not females, suggesting that afferent renal nerves are involved in increase in BP via V1b AVP receptor. We additionally show that despite normal BP, female CMS rats display increased renal sympathetic nerve activity (RSNA; 2.39 ± 0.23 bursts/beat vs. 1.44 ± 0.12 bursts/beat, p < 0.005) measured directly with implanted telemetry in conscious rats over one week and aortic stiffness, as evidenced by increased aortic pulse wave velocity (173.2 ± 50.9 mm/s vs. - 10.7 ± 54.6 mm/s in controls, p = 0.0393).

**NEW & NOTEWORTHY:** We show that renal denervation mitigates the rise in blood pressure (BP) in a model that is not genetic nor diet-dependent, the chronic mild stress model (CMS). Specifically, we demonstrate the role of afferent, rather than efferent, renal nerves in mediating the rise in BP in male CMS rats. Finally, we report that renal sympathetic nerve activity, but not BP, is elevated in female CMS rats measured by telemetry over seven days in conscious rats.

## INTRODUCTION

The kidney is richly innervated with renal sensory (afferent) and sympathetic (efferent) fibers which allows for a two-way communication with the brain nuclei responsible for integration of information that governs the cardiovascular and renal function: blood pressure, renal plasma flow, glomerular filtration rate (GFR) and electrolyte balance. Renal afferent nerves are responsible for mechano- and chemo-sensing, relaying the information from the kidney to the premotor nuclei in the paraventricular nucleus (PVN) of the hypothalamus and the rostral ventrolateral medulla (RVLM) (1). The appropriate responses are then integrated and effected via postganglionic neurons which project to different organ beds: vasculature, heart and kidney (2). That increased renal sympathetic nerve activity (RSNA) elicits the rise in blood pressure (BP) has been shown to occur by at least three mechanisms: 1) increase in reabsorption of sodium and water in the kidney tubules, 2) reduction in renal blood flow (via afferent arteriole constriction) and GFR, and 3) renin release via juxtaglomerular apparatus, activating the renin–angiotensin–aldosterone cascade (RAAS) (1). Therefore, modulation of renal nerves to improve BP control has become a topic of intense investigation over the last 10-15 years, starting with promising proof-of-principle clinical trials (3, 4) and culminating in accumulation of significant evidence supporting renal denervation (RDNx) as a durable strategy for uncontrolled hypertension (5). That is to say, we know RDNx works - we just do not know how exactly it works.

Significant preclinical evidence has amassed supporting RDNx as a successful strategy in several different models of hypertension: spontaneous hypertensive rat (SHR) (6), deoxycorticosteroid acetate (DOCA) - salt (7, 8), Dahl-salt (DahlS) (9), dietary fructose and salt feeding (10), but not angiotensin II (Ang II) infusion rat model (11). However, a significant knowledge gap exists regarding the role of renal nerves, both afferent and efferent, in pathobiology of hypertension given the multifactorial etiology of the disease beyond dietary salt and genetics. Although salt-sensitivity plays a significant role in BP regulation and hypertension development, other lifestyle factors, such as psychosocial stress, can contribute to elevation of BP. Chronic stress that leads to depression and other major depressive disorders has been linked to increased risk of cardiovascular morbidity and mortality, according to several major epidemiological studies, such as INTERHEART (12, 13) and INTERSTROKE (14-16). A potential link between stress and autonomic regulation involves evidence that supports the role of centrally acting arginine vasopressin (AVP) in modulating the release of adrenocorticotropic hormone (ACTH) which, in turn, modulates the hypothalamic-pituitary-axis (HPA) response to stress (17, 18). Specifically, the PVN has been shown to provide inputs into the brainstem and spinal cord loci crucial for regulating cardiovascular function (19). We recently showed that intracerebroventricular microinjection of AVP into the PVN of rats exposed to 4-weeks of chronic mild stress (CMS) stimulates renal sympathetic activity leading to increased blood pressure and heart rate (20, 21). The CMS rat has been used to model human chronic stress and depression, recapitulating the behavioral (22, 23) and neurocardiovascular phenotype (20, 24) observed in depressed humans. Although some published work studying CMS from the behavioral perspective used both males and females (25), our published work (20, 26) and that of others (24, 27, 28) studying neurocardiovascular outcomes used male rats only. Our recently published study using combined pharmacological antagonism of AVP receptors, V1a and V1b, in the PVN supports their role in sympathoexcitation in chronic stress, albeit only in male rats (20). However, the role of renal afferent inputs into the central neurocardiovascular nuclei (like the PVN and RVLM) and the effect on vasopressinergic signaling in CMS has not been investigated. Additionally, our recent work showed that female CMS rats remain normotensive, unlike their male counterparts, revealing the existence of sex-specific dichotomy in BP regulation in CMS (29).

The current study was completed to establish the relative contribution of renal afferent and efferent nerves to BP regulation in CMS and to explore potential effects on AVP signaling. Additionally, we set out to describe the sex-specific dichotomy of BP regulation and sympathetic activation in CMS. We hypothesized that: 1) selective ablation of renal afferent nerves (ARDNx) and total removal of all renal nerves (TRDNx) would reduce BP in male CMS rats to the same extent; 2) that ARDNx would alter the expression of AVP receptors, V1a and/or V1b; and 3) that despite normal BP, female CMS rats would display sympathoexcitation to renal and vascular beds by direct measures of RSNA obtained with telemetry in conscious chronically instrumented female rats and aortic pulse wave velocity, respectively.

## MATERIALS AND METHODS

Studies described in this manuscript were completed in male (n = 23) and female (n = 12) Sprague-Dawley rats, weighing 200-225 g (9-11 weeks old) purchased from Charles River Laboratories (Raleigh, NC). Upon receipt from the vendor, rats were housed under controlled conditions (lights on 7AM-7PM, 21-23^°^ C) for at least 48 hours to acclimatize, with *ad libitum* access to water and normal rat chow (Teklad Global 2018). All procedures were carried out in accordance with the principles of the National Institutes of Health’s *Guide for the Care and Use of Laboratory Animals*. All procedures and protocol were approved by the Wayne State University Institutional Animal Care and Use Committee (protocol # 23-05-5867).

### Group Assignments

Female rats were assigned to either the control group or chronic mild stress (CMS) group, at random (n = 6 each). The control group was housed under normal, control conditions for 4 weeks. The CMS group was exposed to the CMS paradigm, as we published before (20, 26, 29). All male rats were exposed to CMS. Briefly, the protocol consisted of exposure of rats to a 7-day schedule of mild stressors (i.e. strobe light, white noise, social housing change from paired to unpaired, constant overnight lighting etc.) repeated for 4 weeks. All surgical procedures (except renal denervation) and protocols described below commenced after the 4-week mark of CMS / control condition in CMS and control groups, respectively. Renal denervation completed in a subset of male animals was done prior to the start of the CMS protocol.

### Surgical Procedures

#### 1. Hemodynamic and RSNA Telemetry

Male rats were instrumented with the BP telemeter (HD-S10, DSI, St. Paul, MN) and female rats with the dual BP and sympathetic nerve activity (SNA) telemeter (TRM56SP, Kaha Sciences Ltd., Auckland, New Zealand). The surgical procedures for BP only telemeter was completed under isoflurane anesthesia (2-3%) via the right femoral artery approach. The inguinal incision was made (∼ 2cm) and the gel-filled catheter was advanced into the abdominal aorta via the right femoral artery while occluding the proximal end. The telemeter body was housed subcutaneously in a small size pocket, and the incision was closed with surgical staples. Analgesia was provided with Ethiqa XR (0.65 mg/kg SQ).

The surgical procedure for dual BP and SNA telemeter was completed under isoflurane anesthesia (2-3%). The abdominal incision (∼ 3-4 cm long) was made to allow housing of the telemeter body in the abdominal cavity, secured to the abdominal wall with suture ribs on the device. The Millar catheter was introduced into the aorta via the femoral artery, while occluding the proximal end. The RSNA electrodes were tunneled subcutaneously to the left flank and extruded via the flank incision. The abdominal incision was closed and with the rat position on its right side, the left renal nerves were identified and dissected from the surrounding tissue via the retroperitoneal approach. The wires were placed under the renal nerve bundle and secured with a small amount of elastomer gel (Kwik-Sil, WPI, Sarasota, FL). The ground electrode was secured to the nearby muscle. The left flank was sutured with absorbable suture, and the skin was closed with surgical staples. The rats were housed in their home cages, fully ambulatory, for 24-hour hemodynamic and RSNA measurement.

#### 2. Renal Denervation

In afferent denervated animals (ARDNx), the left kidney was subjected to afferent-only renal denervation while the right kidney was subjected to total renal denervation. In total renal denervated animals (TRDNx), both the left and the right kidney were subjected to complete removal of all renal nerve bundles. The denervation procedures were completed as previously described (8). Briefly, ARDNx was completed by exposing the kidney via retroperitoneal approach and periaxonal application of capsaicin (33 mM), while TRDNx was achieved by surgical removal of all renal nerves followed by periaxonal application of 10% phenol in ethanol. Sham procedure was completed by following the same steps, without the manipulation of renal nerves. The incisions were closed with absorbable suture (muscle layer) and surgical staples (skin). Rats were given Ethiqa XR (0.65 mg/kg SQ) and meloxicam (1 mg/kg SQ) for analgesia and no antibiotics were administered.

Confirmation of ARDNx and TRDNx efficiency was completed with assessment of calcitonin gene-related peptide (CGRP) via ELISA and norepinephrine (NE) levels via high-performance liquid chromatography, respectively, as we reported before (8).

### Experimental Protocols

Measurements of all hemodynamic parameters and RSNA were completed in conscious, unanesthetized, freely moving rats in their home cages. Pulse wave doppler was completed under isoflurane anesthesia.

### Experiment 1: Effect of ARDNx and TRDNx on hemodynamic parameters in male CMS rats

Twenty-three male Sprague Dawley rats were randomly assigned to either the SHAM (n = 7), ARDNx (n = 8) or TRDNx group (n = 8). The rats recovered for 3-4 days and were then subjected to the CMS protocol, as described above, for 4 weeks. During the last week, they were instrumented with BP telemeters and allowed to recover for 2-3 days. Hemodynamic parameters were measured continuously with Ponemah software (Data Sciences International, St Paul, MN) and the data represent 24-hour averages after 4 weeks of CMS.

### Experiment 2: Hemodynamic, RSNA and aortic pulse wave velocity (aPWV) measurements in female CMS rats

Twelve female Sprague Dawley rats were randomly assigned to either CMS (n = 6) or control (n = 6) condition for 4 weeks. In the last week, they were instrumented with dual BP and SNA telemeters and allowed to recover for 2-3 days before hemodynamic and renal sympathetic measurements were obtained continuously for 7 days. The data represent 24-hour averages on Day 1 (i.e. immediately following the 4-week mark) and Day 7 (7 days later). All parameters were obtained with LabChart 8 (ADInstruments, Colorado Springs, CO) using sampling frequency of 2-minute intervals equally spaced 4 times per hour. Hemodynamic parameters were analyzed with blood pressure module (i.e. BP, HR, PP). Renal sympathetic activity was obtained by first applying calibration parameters according to the manufacturer’s instructions. The signal was then filtered using a band pass filter 200-3,000 Hz. Renal sympathetic nerve bursts were identified automatically using digital peak detection software (LabChart 8, ADInstruments). Burst duration profiles were quantified using the full width at half-maximum amplitude (width 50) to determine the temporal characteristics of individual sympathetic discharges. Bursts were counted and divided by the heart rate to quantify RSNA. The decay time constant, Tau (τ), represents the time required for the falling phase of a detected burst (i.e. sympathetic burst washout period). The burst recovery phase was quantified by calculating Tau, determined via a mono-exponential fit applied to the falling phase of each individual burst from 90% to 5% of peak amplitude. On the last day, aortic pulse wave doppler was completed in n = 5 rats in each group. The rats were anesthetized with isoflurane (2-3%) and pulse wave doppler of the ascending aorta was completed to obtain aortic pulse wave velocity (aPWV) in control and CMS females, as we reported before (30), using Vevo2100 ultrasound.

### Experiment 3: RNA extraction and RT-qPCR of vasopressin receptor expression

In a subset of animals (CMS and ARDNx males and CMS females), upon euthanasia, bilateral PVN tissue was collected with a sample corer, 15G (Fine Science Tools, Foster City, CA) and stored in RNA later (Thermo Fisher Scientific, Waltham, MA) at -80^°^C. Extraction of RNA was completed with RNeasy MiniKit (QIAGEN, Germantown, MD) and quantitated using NanoDrop (Thermo Fisher Scientific, Waltham, MA). Quantification of vasopressin receptor expression was completed with one step RT-qPCR using iTaq Universal SYBR Green one step kit (BioRad, Hercules, CA) and primers as follows: RQP214997-Rat Avpr1b (NM_001289800.2) and RQP051521 – Rat Avpr1a (NM_053019.2) (GeneCopoeia, Rockville, MD). Normalization was completed with GAPDH using primer set RQP049537 - Rat Gapdh (NM_017008.4) (GeneCopoeia, Rockville, MD). One-step RT-qPCR thermocycling reaction was completed using ABI 7700 instrument (Applied Biosystems Inc., Forest City, CA).

### Statistical Analyses

All non-repeated measures were analyzed with one-way ANOVA with Tukey’s multiple comparisons test. Comparisons between parameters in which only two groups exist (i.e. data for control vs CMS females), unpaired t-test with Welch’s correction was used. For repeated measures of 24-hour RSNA data, two-way ANOVA (or a mixed-model) with Sidak post-hoc test was used, not assuming sphericity. Statistical power was re-assessed post-hoc to assure that it is maintained for all parameters at 0.8 at the minimum, with significance level set to p < 0.05. All data are presented as mean ± SEM.

## RESULTS

### Experiment 1: The Role of Renal Afferent Nerves in Blood Pressure Regulation in Male CMS Rats

Both ARDNx and TRDNx reduced mean arterial pressure (MAP) after 4 weeks of CMS exposure in male rats. MAP was similar in ARDNx (104 ± 2 mmHg, p = 0.003) and TRDNx (101 ± 3 mmHg, p < 0.001) compared to SHAM (118 ± 1 mmHg), while there were no differences between ARDNx and TRDNx (Fig. 1A). Additionally, both MAP components, systolic blood pressure (SBP) and diastolic blood pressure (DBP), followed the same trend. SBP was lower in ARDNx (128 ± 3 mmHg, p = 0.0095) and TRDNx (124 ± 3 mmHg, p = 0.008) compared to SHAM (142 ± 1 mmHg) (Fig. 1B). Likewise, DBP was lower in ARDNx (87 ± 2 mmHg, p = 0.0392) and in TRDNx (86 ± 5, mmHg, p = 0.0264) compared to SHAM (99 ± 2 mmHg), while there were no differences between ARDNx and TRDNx for neither SBP nor DBP (Fig. 1C). ARDNx and TRDNx caused no change in heart rate (Fig. 1 D) nor pulse pressure (Fig. 1E).

**Figure 1.**
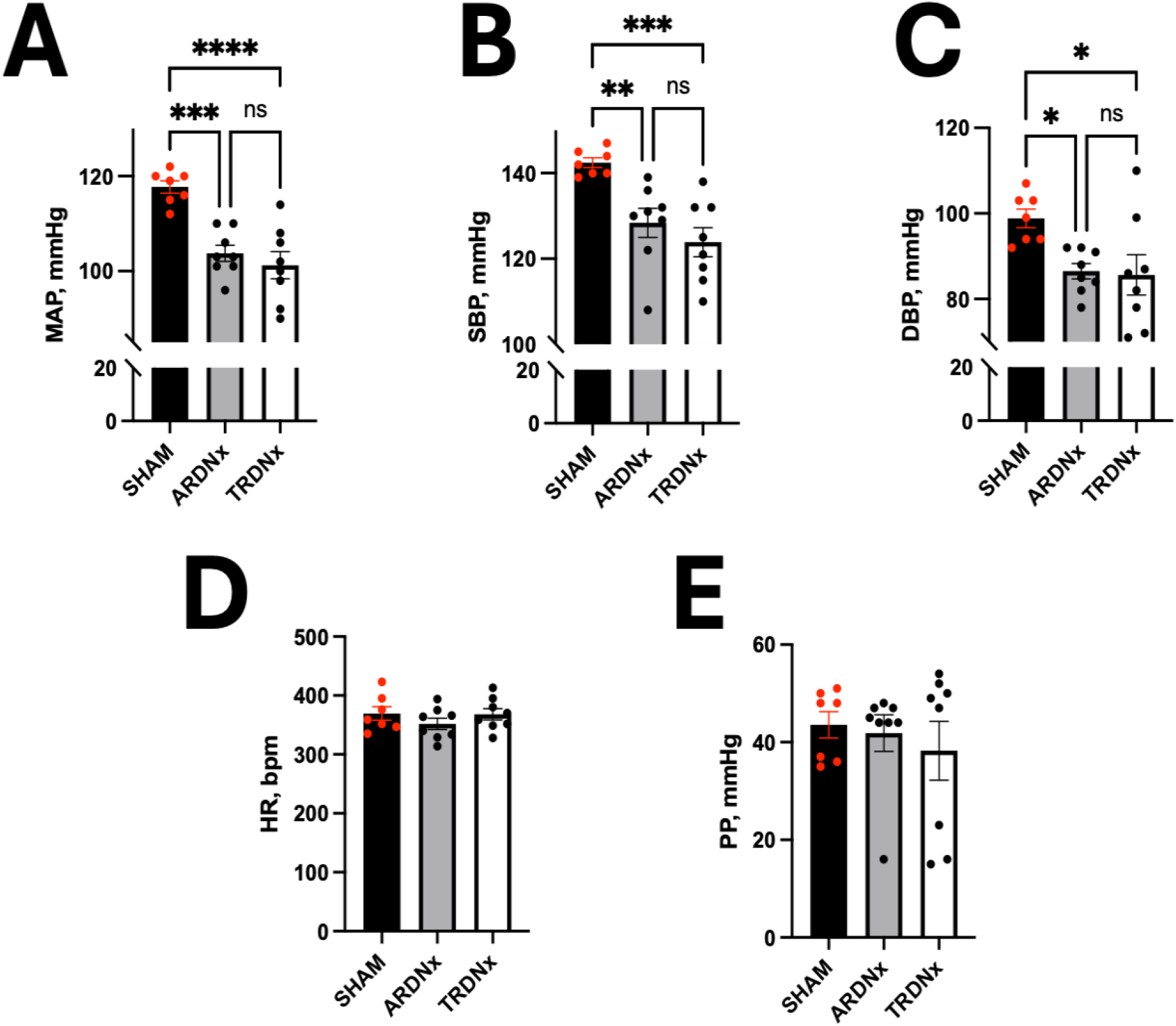
Afferent renal denervation (ARDNx) and total renal denervation (TRDNx) reduce blood pressure equally in CMS male rats. ARDNx and TRDNx reduce mean arterial pressure (MAP, panel A), and both of its components, systolic (SBP, panel B) and diastolic blood pressure (DBP, panel C), after 4-week exposure to chronic mild stress (CMS) in male rats, while having to effect on heart rate (HR, panel D) or pulse pressure (PP, panel E). Data are shown as mean ± SEM. * *P* < 0.05, ** *P* < 0.01, *** *P* < 0.005, **** *P* < 0.001 vs. SHAM. One-way ANOVA with Tukey’s post-hoc test. SHAM: n = 7, ARDNx: n = 8, TRDNx: n = 8.

#### Denervation efficacy

Denervation efficacy for TRDNx and ARDNx was determined with norepinephrine (NE) and calcitonin gene-related peptide (CGRP) kidney content, respectively. NE content was reduced significantly by TRDNx (15.94 ± 9.48 μg / g of renal tissue) compared to SHAM (86.39 ± 7.56 μg / g of renal tissue, p = 0.002) and ARDNx (86.69 ± 13.48 μg / g of renal tissue, p = 0.0009), indicating a successful removal of all renal nerves, while preserving the sympathetic fibers in ARDNx (Fig 2A). Ablation of afferent renal nerves was successful in both ARDNx and TRDNx (Fig. 2B), as indicated by reduced CGRP content in both (12.38 ± 1.06 and 18.44 ± 2.26 pg/mg of renal tissue, respectively) compared to SHAM (36.81 ± 5.03 pg /mg of renal tissue, p = 0.0002 and p = 0.0031 vs. ARDNx and TRDNx, respectively).

**Figure 2.**
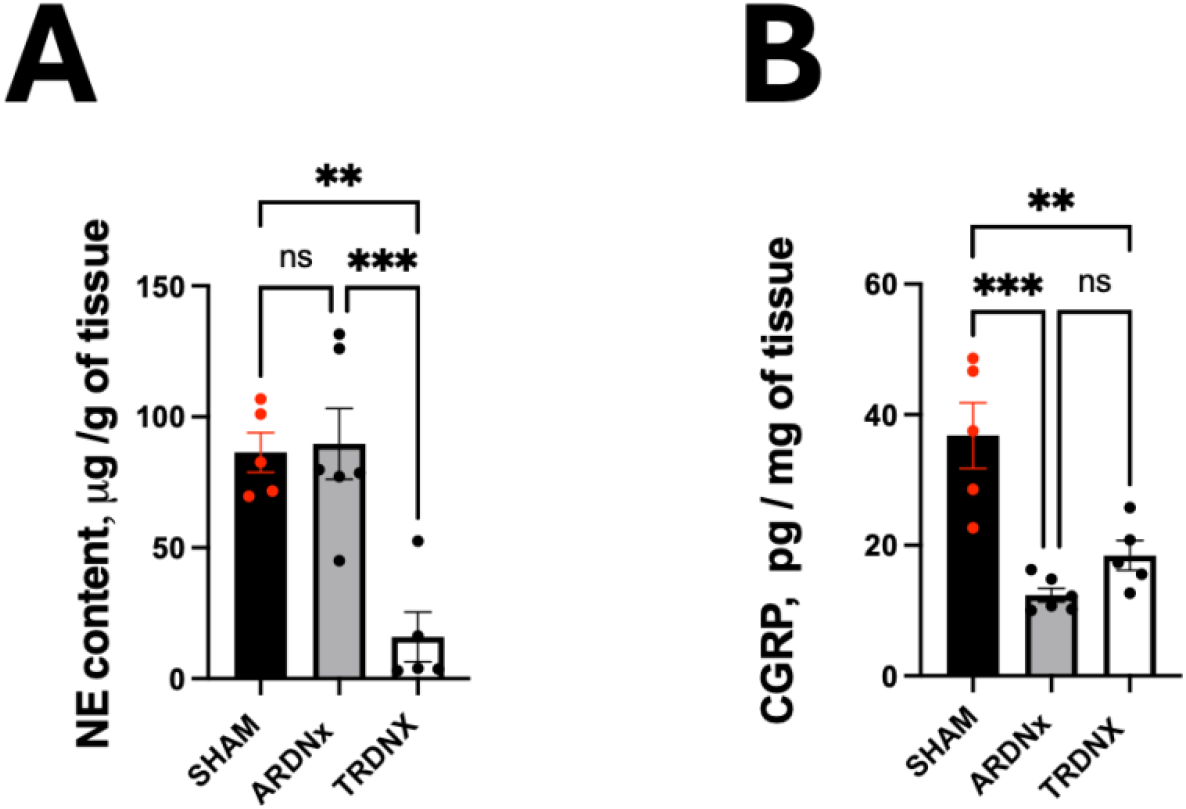
Denervation efficacy in afferent renal denervation (ARDNx) and total renal denervation (TRDNx). Norepinephrine (NE) content was reduced only in TRDNx (A) indicating successful removal of renal sympathetic nerves, which were preserved in ARDNx. Likewise, successful removal of afferent nerves is indicated by reduced calcitonin gene-related peptide (CGRP) in both ARDNx and TRDNx (B). Data shown as mean ± SEM. ** *P* < 0.01, *** *P* < 0.005, **** *P* < 0.001 vs. SHAM. One-way ANOVA with Tukey’s post-hoc test. SHAM: n = 5, ARDNx: n = 6, TRDNx: n = 5.

### Experiment 2: Female CMS Rats are Normotensive but Have Increased Renal Sympathetic Nerve Activity

Unlike male CMS rats, female Sprague Dawley CMS rats remained normotensive after 4 weeks of CMS (Table 1). However, renal sympathetic nerve activity (RSNA) was higher despite the maintenance of normal BP in CMS females (Fig. 3). Specifically, burst incidence was increased after 4 weeks of CMS (2.39 ± 0.23 bursts/beat) compared to control (1.44 ± 0.12 bursts/beat, p = 0.0224, Fig. 4A and Table 2), and it remained consistently increased after 7 days (2.18 ± 0.17 vs. 1.52 ± 0.12 bursts/beat, Figs. 4A and 4B, p = 0.0310). There was no difference in change of RSNA incidence over time between controls and CMS female rats. Over the seven days following CMS, tau was shorter in CMS rats (8.410 ± 0.719 ms) compared to controls (11.874 ± 0.879 ms, p = 0.049, Fig. 4C) on day 1. However, statistical significance was lost seven days later (9.917 ± 1.303 ms vs 11.356 ± 0.905, p =0.054, Table 2).

**Table 1.** Hemodynamic parameters in female control and CMS Sprague Dawley rats.

|  | CONTROL (n = 6) | CMS (n = 6) |
| --- | --- | --- |
| MAP (mmHg) | $98 \pm 4$ | $88 \pm 4$ |
| SBP (mmHg) | $123 \pm 4$ | $117 \pm 4$ |
| DBP (mmHg) | $81 \pm 5$ | $67 \pm 4$ |
| HR (bpm) | $436 \pm 17$ | $395 \pm 9$ |
| PP (mmHg) | $42 \pm 4$ | $48 \pm 1$ |

Data are presented as means ± SEM.; n, number of rats; MAP, mean arterial pressure; SBP , systolic blood pressure; DBP , diastolic blood pressure ; HR, heart rate ; PP, pulse pressure. No statistical significance found in any of the parameters between control and CMS groups.

**Table 2.** 24-hour RSNA burst characteristics in female control and CMS Sprague Dawley Rats measured over 7 days post-CMS/control protocol completion.

|  | DAY 1 |  | DAY 7 |  |
| --- | --- | --- | --- | --- |
|  | Control | CMS | Control | CMS |
| **Incidence (bursts/beat)** | 1.44 ± 0.12 | 2.389 ± 0.23 ** | 1.52 ± 0.12 | 2.18 ± 0.17 * |
| **Tau (ms)** | 11.874 ± 0.879 | 8.410 ± 0.719 * | 11.356 ± 0.905 | 9.917 ± 1.303 |
| **Width50 (ms)** | 24.420 ± 1.444 | 18.502 ± 2.320 | 24.016 ± 1.490 | 20.819 ± 1.876 |

Data are presented as means ± SEM. Control = control rats (n=5), CMS = chronic mild stress rats (n = 6). \* $P < 0.05$ , \*\* $P < 0.005$ vs. time-matched Control. Two-way ANOVA with Šídák's multiple comparisons test.

**Figure 3.**
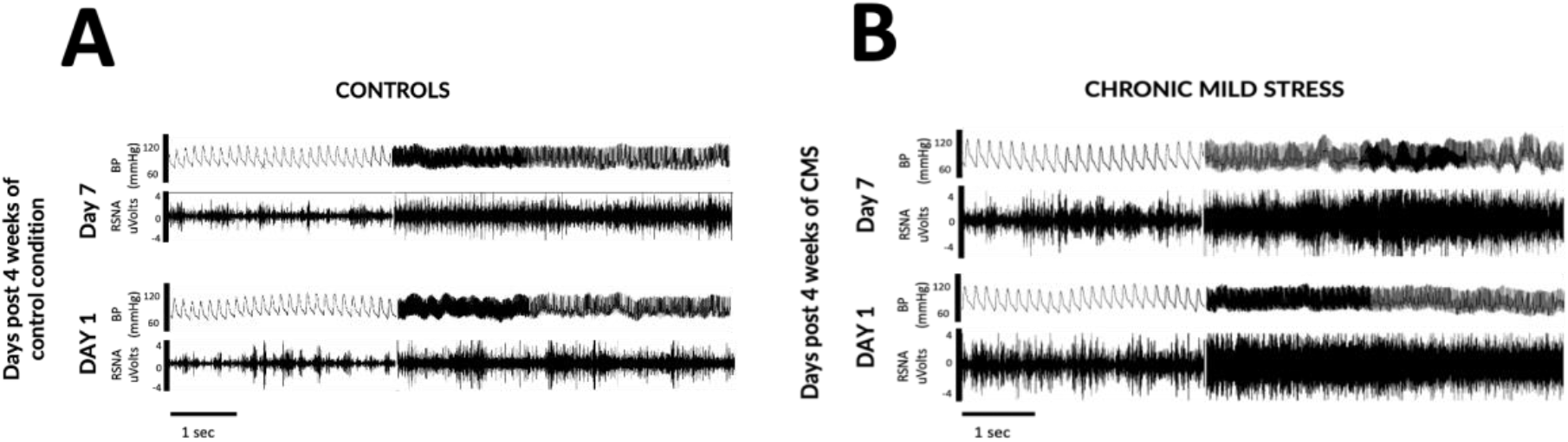
Representative raw blood pressure (BP) and renal sympathetic activity (RSNA) trace after 4 weeks of the control condition (A) and CMS (B). The traces are shown for the same animal within each condition. BP and RSNA were measured with implanted telemetry (dual BP+SNA TRM56S, Kaha) and the data obtained with LabChart. RSNA signal was filtered with band pass of 200-3,000Hz.

**Figure 4.**
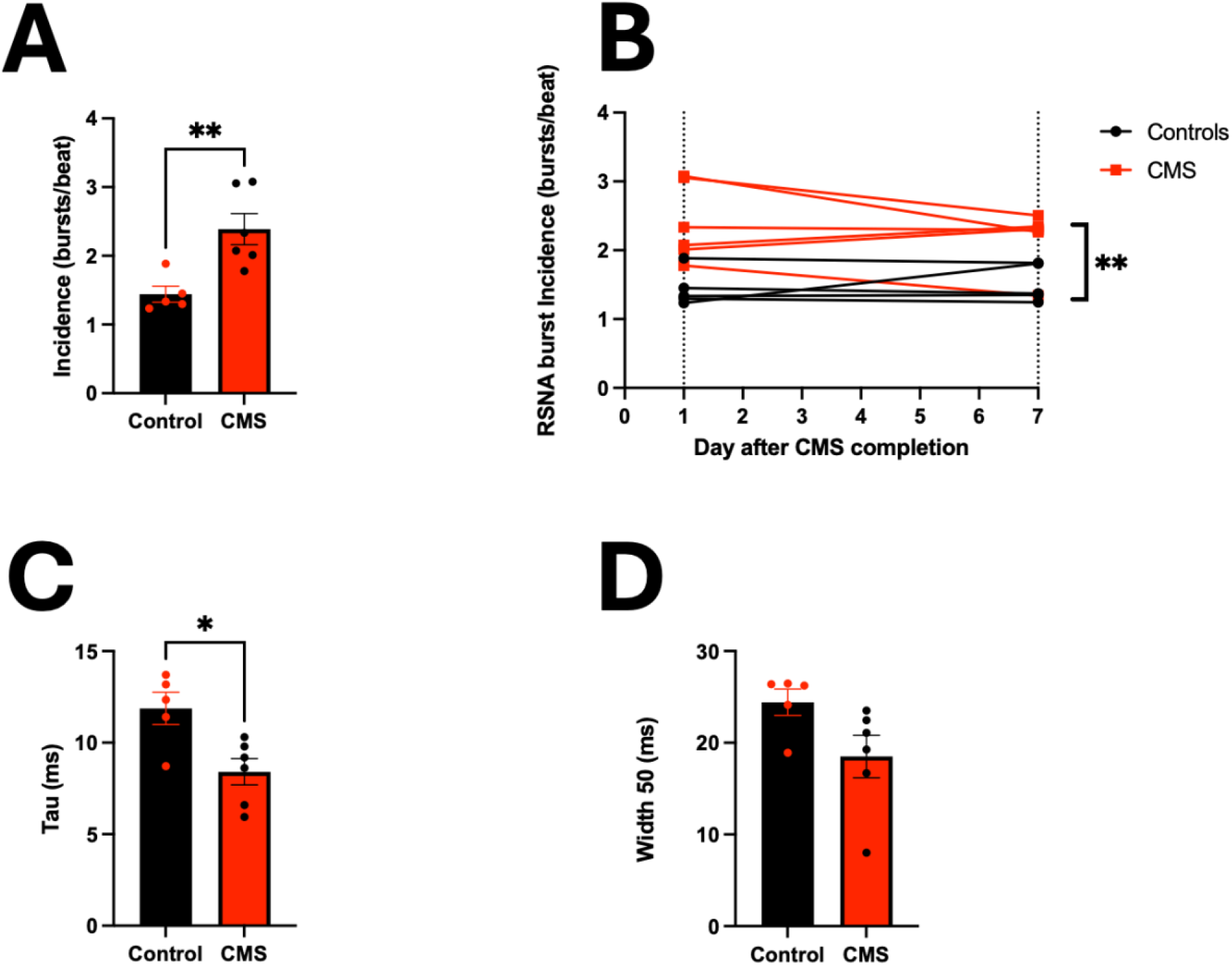
RSNA burst characteristics in control and CMS female Sprague Dawley rats on days 1 and 7 after control/CMS paradigm, respectively. Burst characteristics are shown for day 1 (A) separately and for days 1 and 7 (B). Tau constant was longer in control rats (C) while RSNA bursts tended to be wider in control vs. CMS rats (D). Data are shown as mean ± SEM. * *P* < 0.05, ** *P* < 0.01 vs. Control in panels A and C, Welch’s t-test. ** *P* < 0.005 vs. Controls panel B, Two-way ANOVA with Šídák’s post-hoc test. Control: n = 5, CMS: n = 6.

Aortic pulse wave velocity (aPWV) was assessed with echocardiography using pulse wave doppler before the onset of control/CMS paradigm and 4 weeks after and the difference in aPWV between these two time points is presented in Fig. 5. Aortic PWV increased after 4 weeks of CMS (by 173.2 ± 50.9 mm/s) while it remained unchanged in control rats after 4 weeks of control condition (-10.7 ± 54.6 mm/s, p = 0.0393).

**Figure 5.**
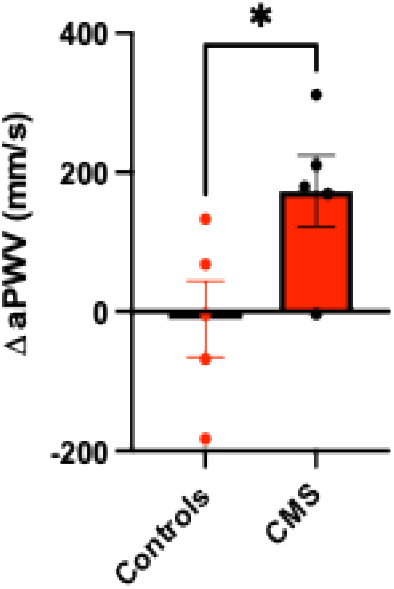
Change in aortic pulse wave velocity (aPWV) measured with echocardiography before and after 4 weeks of control /CMS paradigm in the respective groups of female Sprague Dawley rats. Rats were lightly anesthetized with isoflurane and pulse wave Doppler was used to obtain aPWV in B-mode. * *P* = 0.0393 by Welch’s t-test. N = 5 rats in each group.

### Experiment 3: The Mechanistic Link between Afferent Renal Nerves and Vasopressin Receptor Density in the Paraventricular Nucleus

Vasopressin (AVP) receptor expression was assessed in bilateral paraventricular nucleus (PVN) of the hypothalamus in rats after 4 weeks of CMS in males with intact renal nerves (SHAM males), in males in which selective removal of afferent renal nerves was completed (ARDNx males) and in CMS females with intact renal nerves (SHAM females). While there was no significant difference in the expression of V1a AVP receptor across the groups (Fig. 6B), V1b receptor expression was significantly higher in CMS males with intact renal nerves (Fig. 6A) compared to both ARDNx males (p = 0.0001) and SHAM CMS females (p < 0.0001), while there was no difference between the latter two.

**Figure 6.**
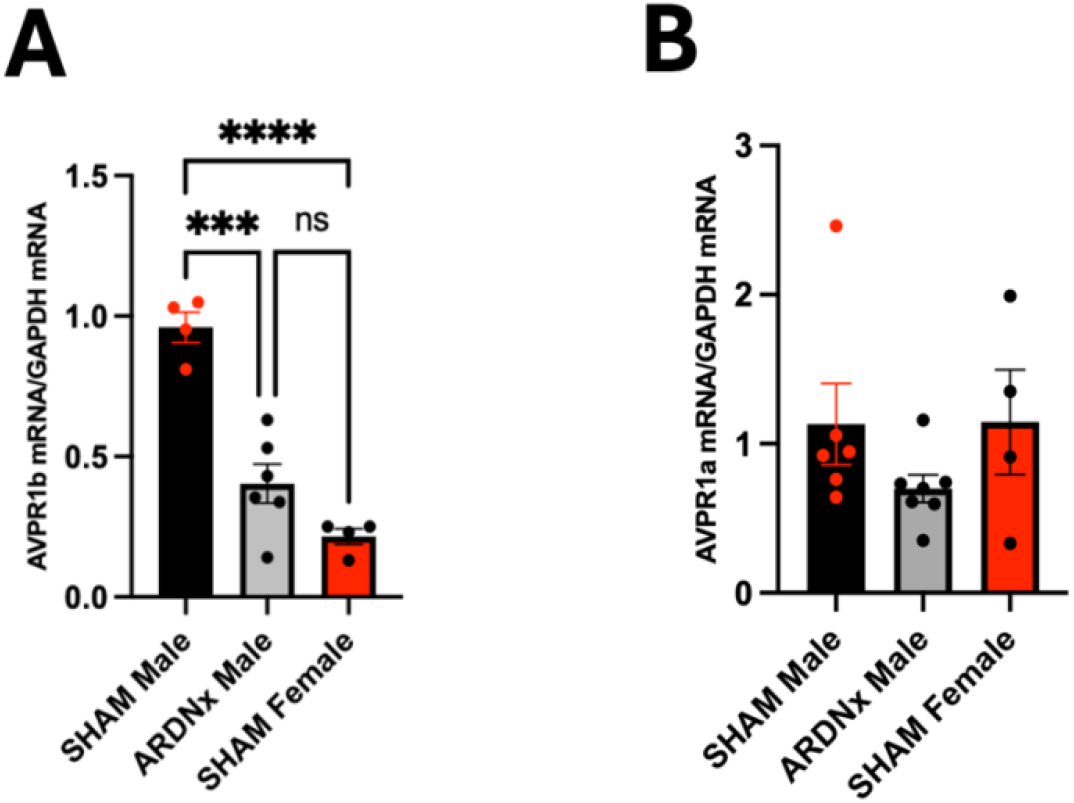
Expression of vasopressin (AVP) receptors in the paraventricular nucleus (PVN) of the hypothalamus. All groups have been exposed to CMS. Bilateral PVNs were collected with tissue corer and RNA was isolated, followed by RT-qPCR using primers specific for V1b (A) and V1a (B) AVP receptors, using GAPDH as normalizing internal control. *** *P* = 0.0001, **** *P* < 0.0001 vs. SHAM male by one-was ANOVA with Tukey’s post-hoc test. SHAM male: n = 6; ARDNx male : n = 7; SHAM female: n = 4.

## DISCUSSION

The objective of this study was to determine the potential role of afferent and efferent renal nerves in regulating BP increase in a rat model of chronic mild stress (CMS) and to demonstrate the sex-specific dichotomy in BP and sympathetic activity regulation. There were four significant findings that we report here for the first time. First, the role of afferent renal nerves is prominent over efferent renal nerves in mediating the rise in BP in CMS male rats. Second, although normotensive (a finding we previously reported (29)), gonadal-hormone intact female CMS rats display increase in RSNA that is evident after 4 weeks of CMS and remains elevated for at least a week afterward. Third, sympathetic activation is not restricted to the renal system, as evidenced by an increase in central vessel stiffness measured with aortic pulse wave velocity (aPWV) after 4 weeks of CMS. Fourth, the increase in BP in CMS is at least partially mediated by afferent renal nerve input into the paraventricular nucleus (PVN) of the hypothalamus which leads to increased vasopressinergic signaling via V1b, rather than V1a AVP receptors.

These findings are consistent with our hypothesis that afferent rather than efferent renal nerves drive the increase in BP in the CMS rat model via increasing vasopressinergic signaling in the PVN and that sympathetic activation may be coordinated in other brain centers and/or may be sex-specific.

### The mechanism of blood pressure increase in chronic mild stress involves afferent renal nerves and AVP V1b receptors in the paraventricular nucleus of the hypothalamus

Renal denervation (RDNx) has been shown to be effective in mitigating hypertension in salt-sensitive (7-9) and genetic animal models (6, 31) but has not yet been applied to a lifestyle-related animal model of hypertension (32, 33). The CMS paradigm has been used to impose a moderate level of stress that roughly equates to the everyday stressors in human life, such as running late for a meeting, flat tire, being stuck in traffic, career stress etc., rather than chronic, more existential stressors. As such, the rise in blood pressure in CMS rat is modest (∼ 7-15 mmHg rise in MAP) (20, 26, 32) compared to some of the more robust hypertension rat models like DOCA, SHR and DahlS (ranging from 50 -100 mHg rise in MAP) (8, 9, 31). Nevertheless, the mechanistic basis of blood pressure rise in any hypertensive milieu merits investigation, given that RDNx has been successfully used to treat human hypertension (5), which is multifactorial in etiology (34, 35).

We previously published in male CMS rats, that the 24-hour MAP rise of ∼7-12 mmHg is accompanied by an increase in RSNA and plasma vasopressin concentration (20, 26). We have also showed that the expression levels of AVP receptors V1a and V1b in the PVN are increased in CMS male rats compared to controls (20). Additionally, most recently we showed that gonadal-hormone intact female Sprague Dawley rats remain normotensive after 4 weeks of CMS (29). In the current study, we aimed to decipher the mechanisms behind these observed phenomena.

We show that both ARDNx and TRDNx reduce MAP equally in male CMS rats, by ∼ 15 mmHg, which is consistent with the magnitude of MAP reduction in other hypertensive rat models, like DOCA (7, 8, 36) and DahlS (9), although TRDNx was only found to reduce MAP in DahlS, and not ARDNx. It is important to note that the DahlS rat used in this study was sourced from a commercial vendor and thus has succumbed to a significant genetic drift from the original colony (37). Therefore, it is unknown whether the DahlS rat from the original colony would recapitulate reduction in MAP with ARDNx. We did not complete ARDNx and TRDNx studies in female CMS rats because they do not become hypertensive after CMS and because it has already been shown that TRDNx reduces blood pressure in normotensive Sprague Dawley rat (7, 36, 38).

Since selective ablation of afferent renal nerves and the ablation of all renal nerves, including the renal sympathetic fibers modulated by the integrating centers like the PVN and RVLM (39), had the same BP reducing effect in CMS males, we speculate that the hypertensive response to CMS is effected by the afferent renal nerves specifically, and as such that the hypertensive response begins by the afferent renal nerve input. Since we showed that vasopressinergic signaling is heightened in CMS male rats (20), we investigated whether selective ablation of renal afferent nerves would impact AVP receptor abundance, in an effort to explain the signal that communicates to the PVN to modulate BP in CMS. The connection between afferent renal nerves and vasopressin has been described over 2 decades ago when it was shown in normotensive Sprague Dawley rats that electrical stimulation of renal afferent nerves triples the plasma vasopressin concentration within 1 hour of stimulation (40). We further describe this interaction between renal afferent nerves and the vasopressinergic system in CMS by showing that reduction of V1b but not V1a receptor abundance in ARDNx CMS males suggests that afferent renal nerves are at least partially responsible for increase in blood pressure via V1b AVP receptor. This is further corroborated by the data observed in the same experiment for females (which remain normotensive after CMS), where there is not an obvious increase in V1b receptor abundance after 4 weeks of CMS (Fig 6). It remains unclear if there are changes in afferent renal nerve activity that underly the blood pressure and molecular effects within the PVN. Future studies are planned to measure this directly.

### Sex-specific dichotomy of blood pressure and sympathetic activity regulation in CMS

Previously published studies explored autonomic dysfunction in chronic stress in rats and indicate the existence of cardiac autonomic dysfunction characterized by an increase in sympathetic frequency domain of heart rate variability (HRV) and a decrease in the parasympathetic component (41, 42), although in one study the power of both sympathetic and parasympathetic frequency domains increased (33). However, these have only been shown in male rats. Although we showed previously that RSNA increases by directly measuring RSNA in conscious CMS rats, this was also only done in males and RSNA was measured only for a short amount of time (∼ 1 hour total) (20, 26). Therefore, we did not measure RSNA in males in this study in order to abide by the 3R principle (replacement, reduction, refinement) in biomedical research. In the current study, we show for the first time that RSNA is increased in female CMS rats despite maintenance of normotension and we do so by obtaining RSNA measurements continuously, over 7 days, by implanted telemetry. This is to our knowledge the first report to have generated data using implanted telemetry to measure 24-hour RSNA concurrently with BP in rats, thereby highlighting the technological innovation aspect of this study. We report that 24-hour averages of RSNA incidence are increased in female CMS rats, which is evident immediately upon completion of the 4-week exposure to the stressors, and is sustained over the following 7 days (Fig. 4). This increase in the number of sympathetic nerve bursts is accompanied by a reduction in burst width and the Tau constant. This evaluation of the RSNA burst kinetics revealed that the burst decay time constant, Tau, was lower in control vs. the CMS female rats (11.874 ± 0.879 ms vs. 8.410 ± 0.719 ms, p < 0.05), indicating a shortening of sympathetic discharge kinetics and a faster return to baseline nerve activity. Similarly, albeit in humans of both male and female sex with chronic anxiety rather than frank depression caused by chronic stress, a previous study reported that muscle sympathetic nerve activity (MSNA) burst amplitude increases compared to non-anxious humans, with no changes in burst frequency (43). We identified no change in burst amplitude, which may be a reflection of sample size as burst amplitude is quite variable for RSNA in rats. Additionally, renal and muscle SNA represent different sympathetic beds and thus may be differentially modulated in physiological stress conditions of varying severity.

The finding that RSNA increases in a setting of normotension in CMS females is novel but the notion that sympathetic activity can increase in normotensive females has been reported before in different physiological scenarios. For example, it has been shown that during mental stress, MSNA responses increased similarly between men and women, but women showed lower pressor effects/ more vasodilation compared to men (44, 45). Additionally, there is a complete lack of relationship between MSNA and BP in healthy young women: women with high MSNA had BP similar to those with low MSNA (46). Although it has been suggested that female sex hormones are responsible for changing sympathetic output as evidenced by MSNA changes throughout the menstrual cycle (47) and with oral contraceptive use (48), the relationship between estradiol, progesterone and sympathetic output remains incompletely understood. Another factor that has been suggested to have a higher potential impact than sex hormones is the difference in subpopulations of adrenergic receptors on the vascular smooth muscle between men and women, with women demonstrating greater β-adrenergic vasodilation potential than men (49). Specifically, women showed less vasoconstriction for the same amount of norepinephrine compared to men. Therefore, there appears to be a greater amount of β-adrenergic vasodilation in women capable of offsetting sympathetically - mediated vasoconstriction. This finding was further corroborated by studies in which a β-blockade with propranolol showed that MSNA and total peripheral resistance increased in tandem in women, and that such relationship was not found to exist in the absence of systemic β-blockade (50). Therefore, gonadal hormone intact CMS female rats could maintain normal BP in light of heightened RSNA via β-adrenergic vasodilation and it would be interesting to see whether this protection could be reversed with the systemic β-blockade.

### Perspectives and Significance

Our study adds on to the existing body of literature by describing an application of renal denervation to models of elevated BP independent of salt sensitivity and volume expansion. We recognize that our study design is a prevention paradigm of RDNx and as such provides insights into the pathogenesis of renal nerves in elevation of BP in chronic stress rather than a reversal paradigm, which is more clinically relevant. Nevertheless, based on current evidence for application of RDNx to humans to improve BP control, additional preclinical studies are needed to understand the contribution of afferent and efferent renal nerve fibers. Coupled with the fact that there currently does not exist a diagnostic test to evaluate the potential efficacy of RDNx to each hypertension case, there is a need for more preclinical studies to characterize the respective contribution of afferent and efferent nerves in the development and treatment of hypertension with multifaceted etiology that can be translated to the clinical scenarios.

Under normal, healthy conditions in rats (but not dogs), BP homeostasis is achieved between the two kidneys via a communication pathway known as the renorenal reflex: increased pressure sensing in one kidney causes the decrease in sympathetic output to the other, prompting the contralateral kidney to excrete more salt and water (51). This reflex may become aberrant in hypertensive conditions where overactivation of afferent renal nerves does not supress, but rather further activates the efferents on the contralateral side, leading to sympathoexcitation (52). Although in our model of ARDNx in addition to all afferent fiber, efferent fibers were also severed on the right side, this does not diminish our findings regarding the role of afferent renal nerves in regulating BP in CMS for at least 2 reasons: 1) ablation of afferent renal fibers on the left side interrupts the renorenal reflex all the same whether the efferent fibers exist on the right side or not and 2) the communication between the kidney and the brain is accomplished with the afferent nerve fibers, which then signal to the BP-integrating centers in the brain to effect the change in BP via mechanisms (i.e. vasopressin release (40)) additional to those that would be effected by the efferent nerve fibers on the right side (i.e. resistance vessel constriction and reduction in renal blood flow (52)). We agree that in experimental paradigms where afferent nerve stimulation is applied (40), the lack of efferent nerves on the contralateral side can be confounding. Additionally, the nerve status in published studies in DOCA rats is identical, where ARDNx model lacks afferents on both sides but also efferents on the right side, because right uninephrectomy is completed first (7, 8, 36). In fact, by leaving the right kidney in and removing all the nerves, our studies argue that some potential factors intrinsic to the kidney do not play a role in BP regulation in CMS, given that both ARDNx and TRDNx groups displayed the same BP reduction magnitude. One limitation to our study is the potential for disrupting parasympathetic innervation of the kidney. The existence of parasympathetic fibers in the kidney has long been disputed, but recent evidence, albeit only in a mouse, shows that in addition to the sympathetic and sensory fibers, the kidney pelvis also harbors parasympathetic nerves (53). Therefore, if such fibers also exist in a rat, they could play a role in BP reduction. It would be expected that the risk of severing parasympathetic nerves (should they exist in a rat) is greater with TRDNx (mechanical severing), and not ARDNx (chemical ablation), and since we saw similar BP reduction magnitude with both TRDNx and ARDNx, the evidence argues against the contribution of parasympathetic fibers to BP lowering effects.

Our finding that ARDNx reduces V1b expression in males argues for a critical role of AVP signaling via V1b in BP regulation in CMS. The finding that V1b expression is similar in CMS females with intact renal nerve fibers and ARDNx males further corroborates this, given that females maintain BP even after 4 weeks of CMS. On the other hand, CMS females display increased RSNA while maintaining normal BP. The sympathetic output is determined by the activity of the central premotor nuclei located in the PVN and RVLM. RVLM neuronal projections extend to pre-ganglionic neurons in the intermediolateral cell column of the spinal cord, which then via postganglionic neurons project to peripheral organs such as heart, arteries, and kidneys (2). Indeed, we provide evidence of such heightened sympathoexcitation in the central vessels via increased aPWV in CMS females. Therefore, it is possible that the heightened activity of the premotor neurons in the RVLM is responsible for the increased RSNA and sympathetic activation to other organ beds (i.e. vascular). Importantly, both RVLM and the PVN activity is modulated by renal mechano- and chemoreceptor reflexes mediated via renal afferent nerves (1), and thus it would be interesting to see whether ARDNx in female CMS rats mitigates central vessel stiffness and RSNA increase.

The significance of our finding that CMS females display increased RSNA even though they maintain normal BP lies in the realization that after menopause, β-adrenergic-mediated protection of BP could potentially disappear. Indeed, it was found in human women that the loss of β-adrenergic vasodilation predominance that is observed in premenopausal women is no longer present after menopause and can therefore explain, at least in part, the rise in hypertension incidence in postmenopausal women (50). In other words, the heightened sympathetic transduction will equate to higher vasoconstriction. In the context of chronic stress, the contribution of increased RSNA to development of hypertension after menopause thus becomes obvious. Perhaps even more clinically relevant is the realization that measuring sympathetic activity is not on the list of routine procedures in the clinic and even in women who are proactive in monitoring their cardiovascular and overall health, measuring BP (which is routinely done) may paint a completely false picture of the CVD risk any given premenopausal woman faces. Such evidence argues for an implementation of additional tests for women, such as non-invasive ultrasound doppler that may reveal vascular function deficits thus allowing for a timelier intervention.

## Notes

### Competing Interest Statement

The authors have declared no competing interest.

